# Effects of a Maternal Ketogenic Diet on Maternal and Offspring Metabolic Health in Mice

**DOI:** 10.64898/2026.07.30.741686

**Authors:** Antonio H. Ruano, Rachel A. Kracaw, Isabella A. Cervantes, Simone Hernandez Ruano, Moises J. Tacam, Kathleen A. Pennington

## Abstract

Ketogenic (keto) diets have gained popularity due to their potential benefits on weight loss and metabolic conditions. However, the consequences of consuming a keto diet before, during, and after pregnancy on maternal and offspring health remain incompletely understood. To investigate this, female Swiss Webster mice were randomly assigned to either a keto or control diet. After 6 weeks on diet, females were mated and maintained on their respective diets throughout pregnancy and lactation. In experiment 1, glucose tolerance tests (GTT) were performed on pregnant dams at gestational day (GD) 16.5. At GD 17.5, dams were euthanized, urine was collected for ketone analysis, and fetal and placental weights were recorded. In experiment 2, dams delivered naturally and one male and one female offspring per dam were retained and weighed weekly. At 12 weeks of age, offspring underwent GTT, followed by euthanasia and serum collection for insulin and leptin analysis. Compared with controls, keto dams exhibited increased body weight, impaired glucose tolerance, elevated urinary ketones, and reduced circulating insulin during late gestation (p<0.05). Fetuses from keto dams were significantly smaller (p<0.05), whereas placental weight and placental efficiency were unchanged. Offspring from keto dams weighed less than controls during early postnatal development but demonstrated catch-up growth by 12 weeks of age. At 12 weeks, keto offspring exhibited a modest but statistically significant improvement in glucose tolerance compared to control offspring (p<0.05). Male keto offspring had significantly decreased serum insulin (p<0.05) compared to male control offspring, while no differences were observed in females. Serum leptin levels did not differ between diet groups, although sex differences present in control offspring were not observed in keto offspring. Together, these findings suggest that maternal ketogenic diet consumption is associated with altered maternal glucose homeostasis during pregnancy and reduced fetal growth in this mouse model. Although offspring exhibited only modest metabolic alterations during early adulthood, the observed changes in growth trajectory and sex-specific metabolic profiles support further investigation into the long-term consequences of maternal ketogenic diet exposure during pregnancy.

## INTRODUCTION

Obesity is a growing global epidemic that contributes to increased health risks and significant economic burden [1]. Obese individuals are at heightened risk for metabolic syndrome, a condition characterized by central obesity, diabetes, hypertension, and dyslipidemia [2]. In the United States, metabolic syndrome accounts for 6-7% of all mortalities [1, 3, 4]. Exposure to an adverse maternal environment during development is a well-established risk factor for later-life metabolic syndrome [2, 3, 5]. This observation was central to the formulation of the ‘developmental origins of health and disease’ (DOHaD) hypothesis, also referred to as fetal programming [3, 6]. Maternal nutrition plays a key role in shaping the intrauterine environment, and numerous epidemiological studies in humans as well as animal models have demonstrated that poor maternal nutrition increases the risk of offspring developing metabolic syndrome, which includes non-alcoholic fatty liver disease (NAFLD), and cardiovascular disease (CVD), later in life [2, 3, 5].

Ketogenic (keto) diets-high in fat, moderate in protein and low in carbohydrates-have gained popularity for their potential benefits in weight loss, metabolic disorders, cardiovascular health, and neurological conditions [7–12]. However, adverse outcomes have also been reported, including increased cholesterol and triglycerides, as well as altered liver lipid metabolism, which are linked to diabetes, CVD, and hypertension [13–15]. More recently, keto diets have been recommended for women with polycystic ovary syndrome (PCOS), a condition affecting 8-13% of reproductive age women [14, 16–18]. Despite the increased consumption of a keto diet consumption in women of reproductive age, the effects of a keto diets during pregnancy and on long-term offspring health remain incompletely understood, with only a limited number of animal and human studies addressing maternal and offspring outcomes [19–23].

Although keto diets have demonstrated metabolic benefits in non-pregnant populations [8, 11, 14, 16], pregnancy represents a unique physiological state characterized by progressive insulin resistance, altered maternal nutrient partitioning, and increased metabolic demands to support fetal growth [24]. Consequently, maternal responses to keto diet consumption during pregnancy may differ substantially from those observed outside of pregnancy, highlighting the need for pregnancy-specific investigation. Animal models provide a valuable opportunity to examine maternal and fetal responses to keto diet exposure under controlled experimental conditions that cannot be readily achieved in humans. However, species-specific differences in energy metabolism and responses to keto diets should be considered when interpreting these findings and translating them to human pregnancy.

Animal studies provide early concerns: in rats gestational keto diet consumption impaired offspring somatic and neurological development [20], while in mice it altered neonatal brain structure and slowed physiologic growth [21, 22]. These alterations in brain structure we subsequently linked to altered offspring behavior [23, 25]. A recent study by Zala et al., demonstrated a ketogenic diet fed during gestation resulted in reduced litter size, altered sex ratio and had sex specific effects on the offspring [19]. However, to date, no studies have examined the impact of maternal keto diet on long-term offspring metabolic health.

The objective of this study was to evaluate the effects of keto diet on maternal and offspring metabolic health. We hypothesized that consumption of a low-carbohydrate, high-fat keto diet before, during and after pregnancy would alter maternal metabolic adaptations during pregnancy and influence offspring metabolic health.. To test this, we evaluated the effects of maternal ketogenic diet consumption on maternal glucose homeostasis during late gestation, fetal growth, and offspring metabolic health using a mouse model.

## MATERIALS AND METHODS

### Animals

The authors assert that all procedures contributing to this work comply with the ethical standards of the relevant national guides on the care and use of laboratory animals (National Institute of Health (NIH) and were approved by the Baylor College of Medicine Institutional Animal Care and Use Committee (IACUC protocol #AN-7439). This study is reported in accordance with ARRIVE guidelines. Figure 1 is a schematic of the animal experimental design which is also described below.

**Figure 1.**
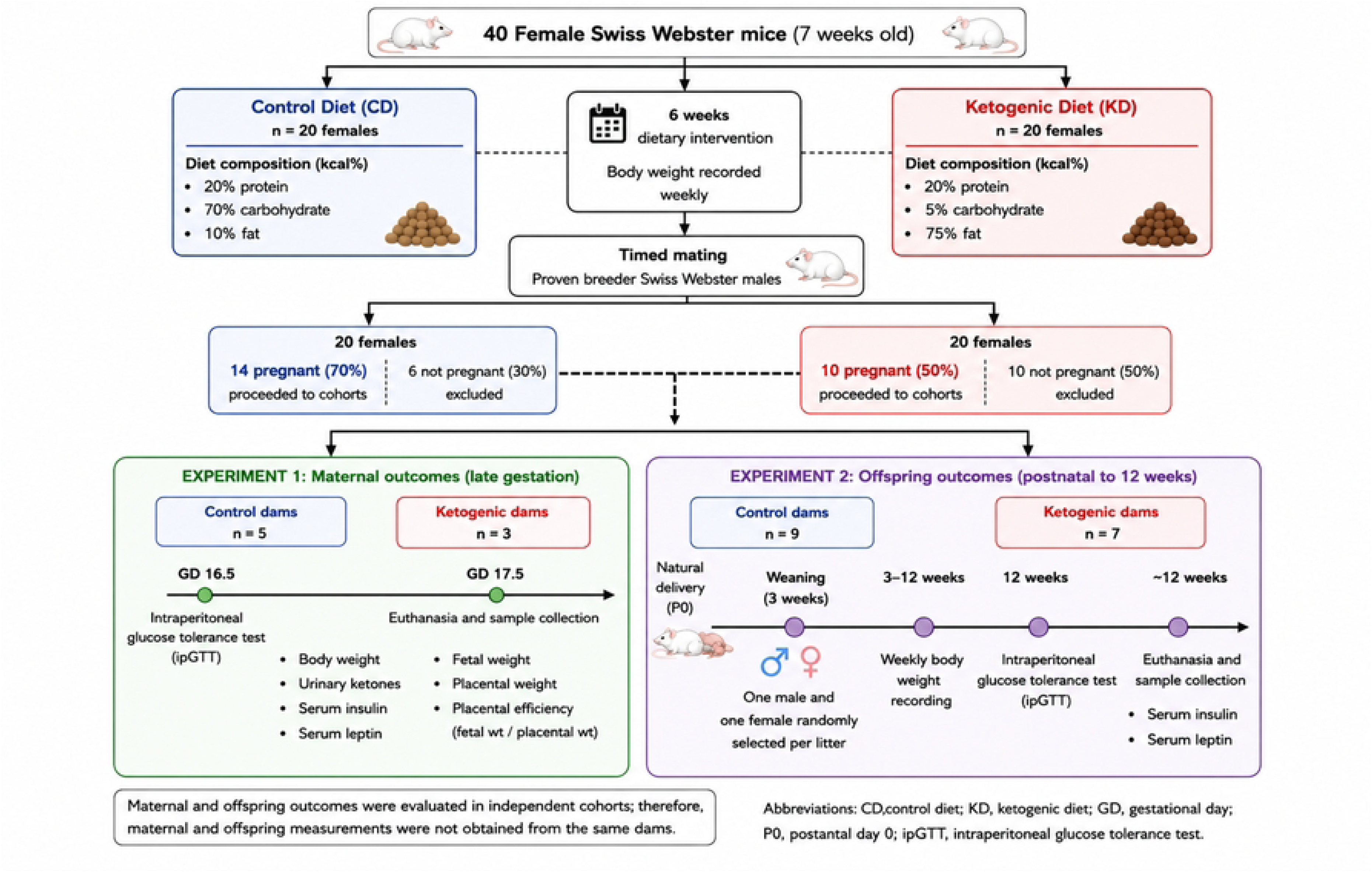
Experimental design and animal allocation for maternal and offspring studies. Forty female Swiss Webster mice (7 weeks of age) were assigned to either a control diet (CD; 20% protein, 70% carbohydrate, 10% fat; *n* = 20) or a ketogenic diet (KD; 20% protein, 5% carbohydrate, 75% fat; *n* = 20) for 6 weeks prior to timed mating with proven breeder males. Pregnancy rates were 70% (14/20) in the CD group and 50% (10/20) in the KD group. Pregnant females were allocated to one of two independent experimental cohorts. **Experiment 1** evaluated maternal metabolic adaptations during late gestation (CD: *n* = 5 dams; KD: *n* = 3 dams). An intraperitoneal glucose tolerance test (ipGTT) was performed on gestational day (GD) 16.5, followed by euthanasia and sample collection on GD17.5 for assessment of maternal body weight, urinary ketones, serum insulin, serum leptin, fetal weight, placental weight, and placental efficiency (fetal weight/placental weight). **Experiment 2** evaluated offspring outcomes from birth through 12 weeks of age (CD: *n* = 9 dams; KD: *n* = 7 dams). Following natural delivery, one male and one female offspring were randomly selected from each litter for longitudinal assessment. Offspring body weight was recorded weekly from 3–12 weeks of age. At 12 weeks, offspring underwent an ipGTT followed by euthanasia and collection of serum for insulin and leptin analyses. Maternal and offspring outcomes were evaluated in independent cohorts; therefore, maternal and offspring measurements were not obtained from the same dams. **Abbreviations:** CD, control diet; KD, ketogenic diet; GD, gestational day; P0, postnatal day 0; ipGTT, intraperitoneal glucose tolerance test.

Six-week-old female Swiss Webster mice were obtained from Charles Rivers Laboratories (Wilmington, Massachusetts, USA) and housed in a 14/10 hour light/dark cycle at 22-24°C with 40-60% humidity. At seven weeks of age, females, weighing 31.5±0.3 grams (see supplementary data files), were randomly divided into two diet groups: control (n=20; 20% protein, 70% carbohydrate, 10% fat; cat#: D12450K, Research Diets, New Brunswick, NJ, USA) or keto (n=20; 20% protein, 5% carbohydrate, 75% fat; cat# D14090501 Research Diets) as previously described [26]. Detailed diet composition, including macronutrient sources and energy density, is provided in Supplementary Table S1. Following 6 weeks of diet exposure, females (approximately 13 weeks of age) were mated to proven breeder males (aged 15-20 weeks) in a ratio of 2 females to 1 male, and the day of copulatory plug was designated as gestational day (GD) 0.5. Following plug detection dams were maintained on their respective diets throughout pregnancy and lactation. Mice were provided ad libitum food and water as well as access to nesting materials.

Because conception rates were lower than anticipated, particularly in the keto diet females, animals were allocated into two independent experimental cohorts to permit completion of both maternal and offspring outcomes. Experiment 1 was designed to evaluate maternal metabolic adaptations during late gestation with n=5, and n=3 keto dams. Experiment 2 was designed to evaluate long-term offspring outcomes with n=9 control and n=7 keto,.for a conception rate of 70% for the control group and 50% in the keto group which was not significantly different between groups. Importantly, these cohorts were independent, and maternal and offspring outcomes were not derived from the same dams, therefore, correlation analysis between maternal and offspring phenotypes were not possible.

Experiment 1: At gestational day (GD) 16.5 glucose tolerance tests (GTT) were performed in control (n=5) and keto (n=3) dams, at GD 17.5 urine was collected by scruffing the dams and applying gentle pressure to the lower abdomen to stimulate urination. Urine was collected into a microcentrifuge tube and then immediately analyzed for urine ketones as described below. Dams were then euthanized via CO^2^ inhalation (30% chamber volume/min) followed immediately by cardiac puncture for blood collection. Blood was collected for future serum analysis and fetal and placental weights were obtained.

Experiment 2: Dams were allowed to deliver naturally. Litter size at birth was recorded; however litters were not standardized or culled. To minimize stress and the risk of pup cannibalism, litters were not disturbed immediately following birth. Consequently, individual pup body weights, maternal body weights during lactation, litter level sex ratios and neonatal survival were not recorded. At weaning, one male and one female offspring from each litter were retained for longitudinal analysis and weighed weekly until 12 weeks of age. In total 9 control and 7 keto dams gave birth, the other females (n=6 control and n= 10 keto) were not pregnant and were euthanized. At 12 weeks of age, GTTs were performed. Offspring were euthanized at 12 weeks of age via CO^2^ inhalation (30% chamber volume/min) followed immediately by cardiac puncture for blood collection. Blood was collected for serum insulin and leptin analysis. For all studies, experimenters were blinded to animal treatment.

### Urine Ketone Analysis

Just prior to euthanasia urine was collected and then immediately applied to ketone test strips (PrimeScreen, Wondofo Biotech, Science City, Huangpu, Guangzhou, China). A visual reading was made and recorded.

### Intraperitoneal Glucose Tolerance Tests

IPGTTs were performed in control and keto dams at GD 16.5 and 12 week old offspring as previously described [27–30] and according to the National Mouse Metabolic Phenotyping Centers protocols for IPGTT (https://www.mmpc.org/shared/document.aspx?id=141&docType=Protocol). Briefly, dams were fasted for 4 hours or offspring were fasted for 6 hours, and then a baseline fasting blood glucose sample was obtained from a venous tail sample. An intraperitoneal injection of glucose (2g/kg) was given. Blood glucose levels were obtained at 15-, 30-, 60- and 120-minutes post glucose injection by removing the tail scab for blood collection. All blood glucose measurements were performed in duplicate using two ReliOn Prime Blood Glucose Monitoring System meters (Walmart, Bentonville, AR, USA).

### Serum Analysis

Serum insulin and leptin were measured in control and keto dams at GD 17.5 and in 12 week old control and keto offspring using a Rat/Mouse Insulin ELISA (EMD Millipore, Billerica, MA, USA) and Mouse Leptin ELISA (EMD Millipore), respectively, as previously described [30, 31] and according to manufacturer’s instructions.

### Statistical Analysis

Statistical analysis was performed using GraphPad Prism (La Jolla, CA, USA). Normality of data was assessed using the Shaprio-Wilk test. When assumptions of normality were not met, equivalent non-parametric analyses (Mann-Whitney) were performed. For maternal ipGTT, data were analyzed using a 2-way repeated measures ANOVA with treatment and time as factors. Area under the curve (AUC) was calculated for ipGTT with units expressed as mg/min/dL. For dam data, AUC, urine keto levels, litter size, fetal and placental weights, serum insulin and leptin a student’s t-test was performed. Fetal and placental weights as well as placental efficiency were averaged per mom, and one value was used per mom for statistical analysis. For all offspring data, to avoid pseudo reduplication and ensure independence of observations, a single male and female offspring per dam were used for analysis, and statistical comparisons were performed using dam as the experimental unit. For offspring body weights and ipGTT a 3-way repeated measures ANOVA was performed with time, sex, and treatment as factors. For offspring AUC, serum insulin and leptin a two-way ANOVA was performed with treatment and sex as factors. To test for outliers for all data the Built-in Outlier test function was utilized in GraphPad Prism using the ROUT method, no outliers were identified and all data is included in the supplementary data files.

## RESULTS

### Consuming a ketogenic diet before and during pregnancy results in altered glucose homeostasis

At GD 16.5, keto dams exhibited impaired glucose tolerance, as indicated by significantly elevated blood glucose levels at 30 and 60 minutes post glucose injection and increased area under the curve (AUC) compared to control dams (p<0.05; Fig. 1A-B). Keto dams also weighed significantly more than control dams at this timepoint (p<0.05, Fig. 1C).

At GD 17.5 keto dams displayed increased adiposity by gross observation, although serum leptin levels were not different between groups (Fig. 1D). Serum insulin levels were significantly decreased in keto dams compared to controls (p<0.05; Fig. 1E). Urinary ketone levels were significantly elevated in keto dams, confirming a state of ketosis (p<0.05; Fig. 1F).

Mean litter size did not differ between control and keto dams at GD 17.5 (Fig. 2A). However, fetus from keto dams had significantly lower mean fetal weights despite similar litter sizes (p<0.05; Fig. 2B), while placental weights and placental efficiency were not different (Fig. 2C-D). These findings indicate that maternal ketogenic diet consumption was associated with altered maternal glucose homeostasis and reduced fetal growth in this mouse model.

**Figure 2:**
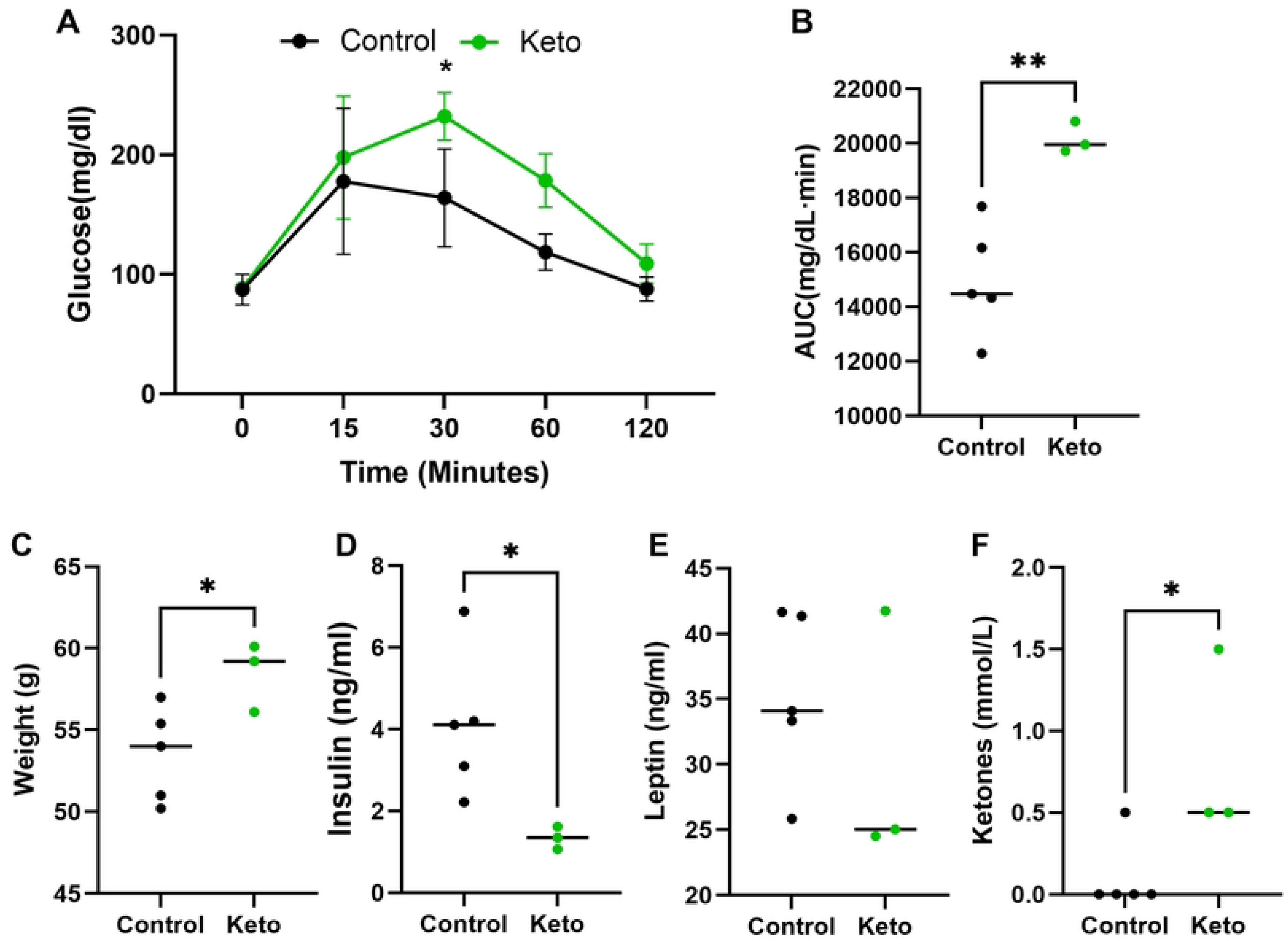
Maternal metabolic outcomes in control and ketogenic diet-fed dams. (A) Blood glucose levels during intraperitoneal glucose tolerance test (IPGTT) at GD 16.5. (B) Area under the curve (AUC) for IPGTT. (C) Maternal body weight at GD 16.5. (D–E) Serum leptin and insulin levels measured at GD 17.5. (F) Urinary ketone levels assessed at GD 17.5. Data are presented as mean ± SE. Statistical analysis was performed using two-way ANOVA for IPGTT and Student’s t-test for other comparisons. Each data point represents one dam (Control n=5, Keto n=3). *p<0.05, **p<0.01.

### Maternal ketogenic diet exposure alters offspring growth and metabolic outcomes

Litter size at birth was not different between control and keto fed-dams (10.4±0.7, 10.9±0.7; p=0.6777). Male and female offspring from keto dams weighed significantly less than control offspring during early postnatal development (p<0.0001; Fig. 3). However, by 12 weeks of age, body weights were not significantly different between groups. Regardless of maternal diet, male offspring weighed significantly more (p<0.0001; Fig. 3).

**Figure 3:**
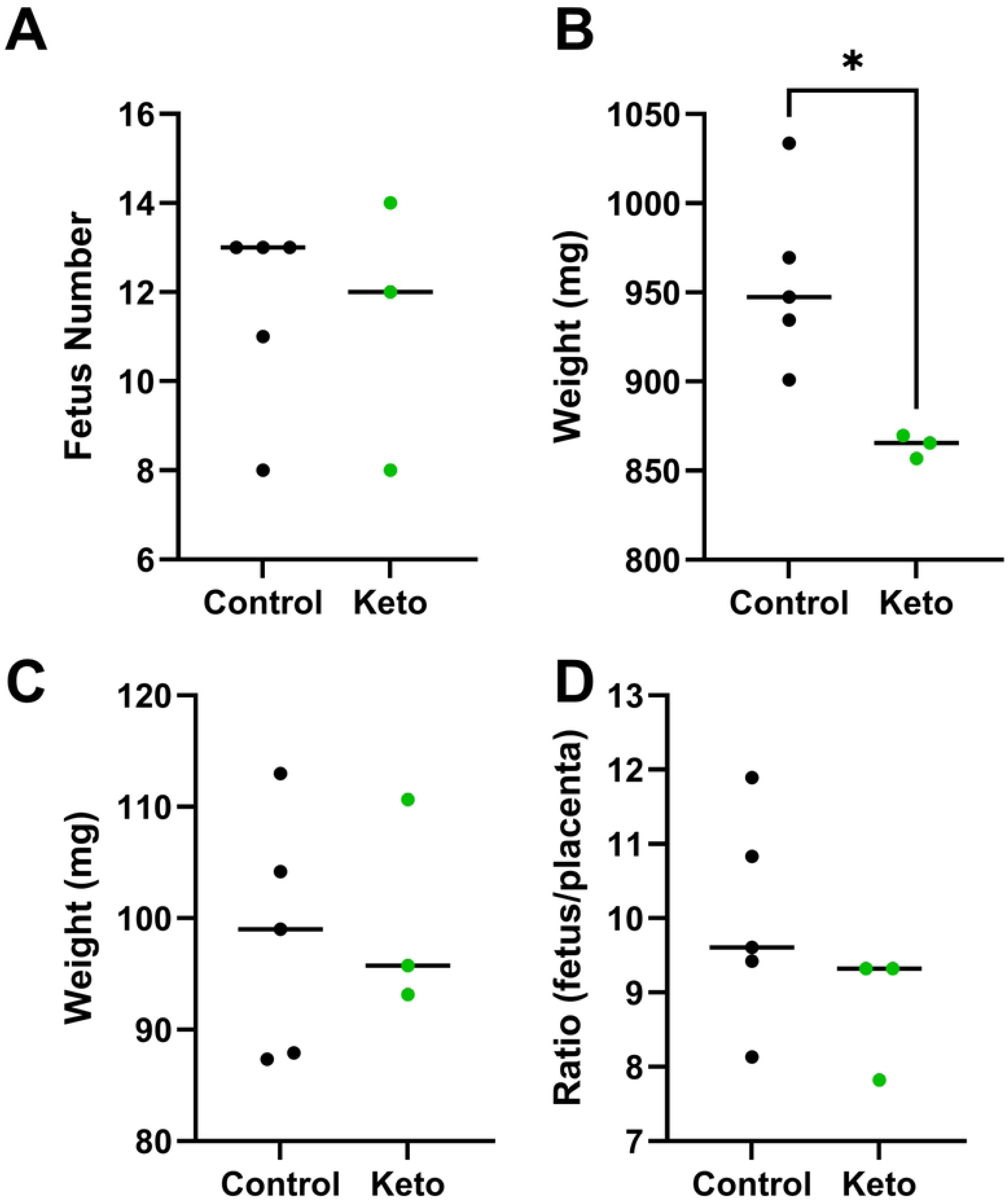
Fetal and placental outcomes at GD 17.5. (A) Litter size per dam. (B) Mean fetal weight per dam. (C) Mean placental weight per dam. (D) Placental efficiency (fetal weight/placental weight). Each data point represents the mean value per dam. Data are presented as mean ± SE. Statistical analysis was performed using Student’s t-test (Control n=5, Keto n=3). *p<0.05.

At 12 weeks of age, glucose tolerance tests revealed that offspring from keto dams exhibited modest but statistically significant improvement in glucose tolerance compared to control offspring (p<0.05, Fig. 4A-B).

**Figure 4:**
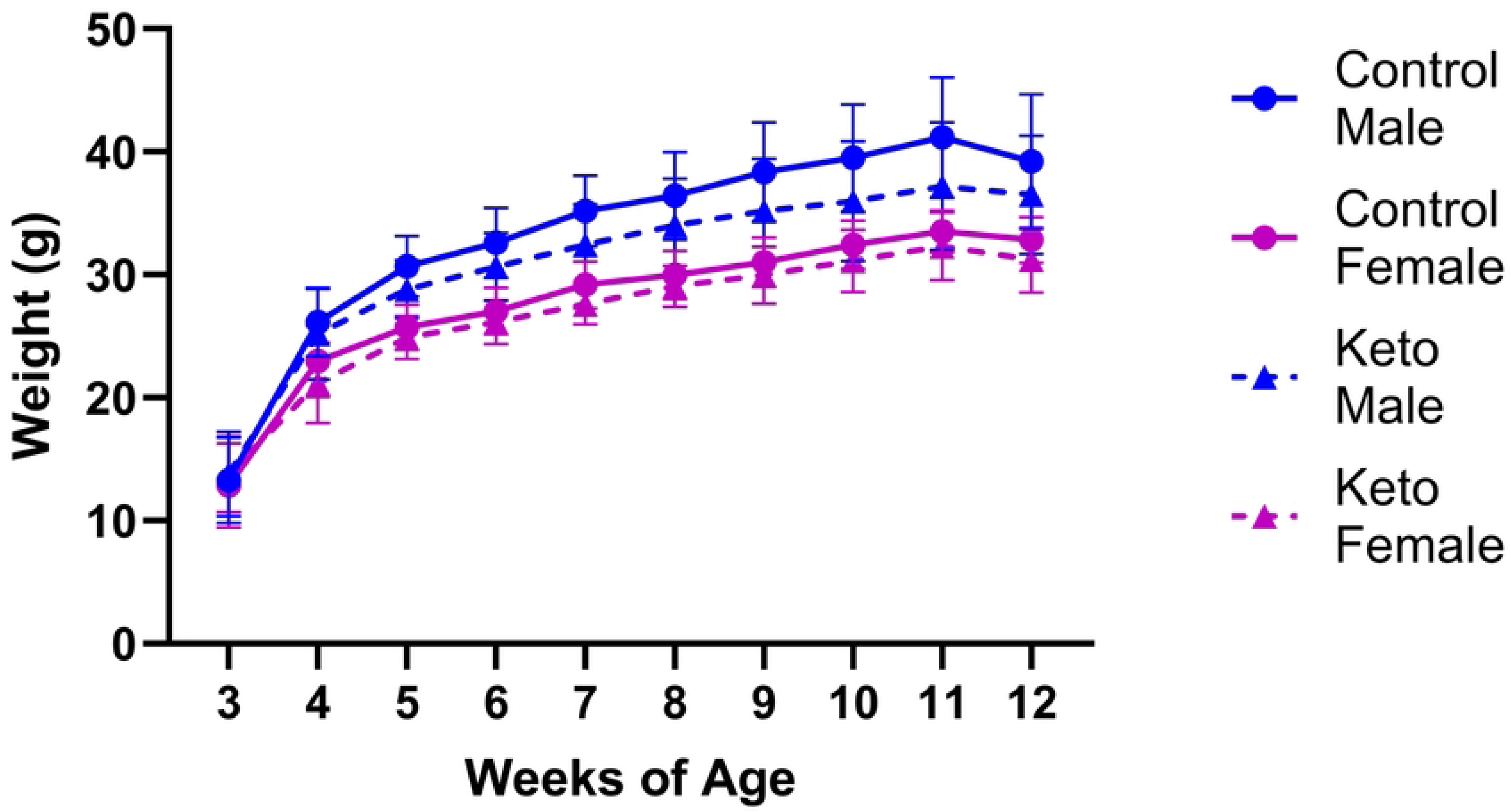
Body weight of offspring from control and ketogenic diet-fed dams. Body weights of male and female offspring measured weekly from weaning to 12 weeks of age. Data represent mean ± SE, with one male and one female per dam included in statistical analyses (Control n=9 dams, Keto n=7 dams). Statistical analysis was performed using three-way ANOVA with time, sex, and maternal diet as factors.

Serum insulin levels were significantly reduced in male keto offspring compared to male controls (p<0.01; Fig 5A), while no differences were observed between female control and keto offspring. Within the control group, male offspring exhibited higher insulin levels than females (p<0.01); however, this sex difference was not observed in keto offspring.

**Figure 5:**
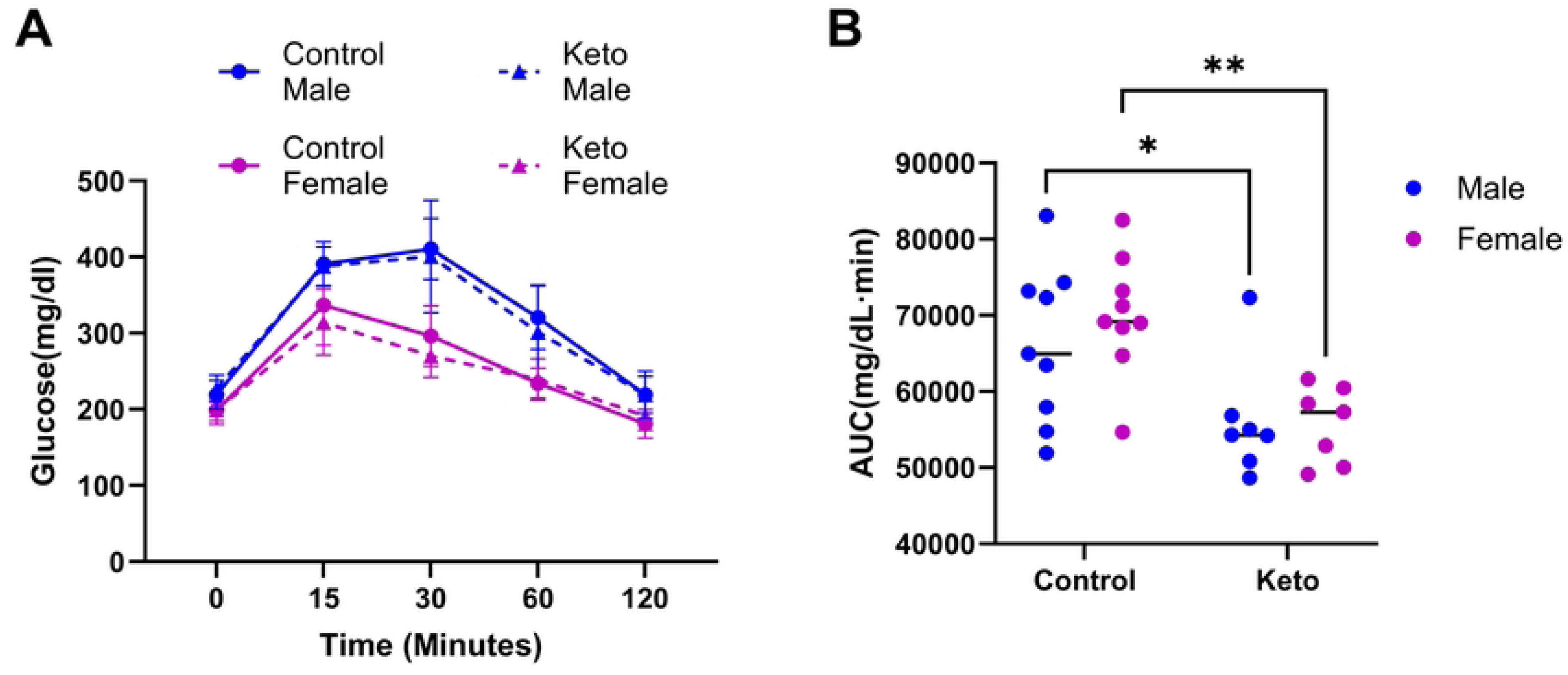
Glucose tolerance in offspring at 12 weeks of age. (A) Blood glucose levels during IPGTT. (B) Area under the curve (AUC). Data represent mean ± SE, with one male and one female per dam included in analysis (Control n=9 dams, Keto n=7 dams). Statistical analysis was performed using three-way ANOVA with sex, maternal diet, and time as factors. *p<0.05, **p<0.01.

**Figure 6:**
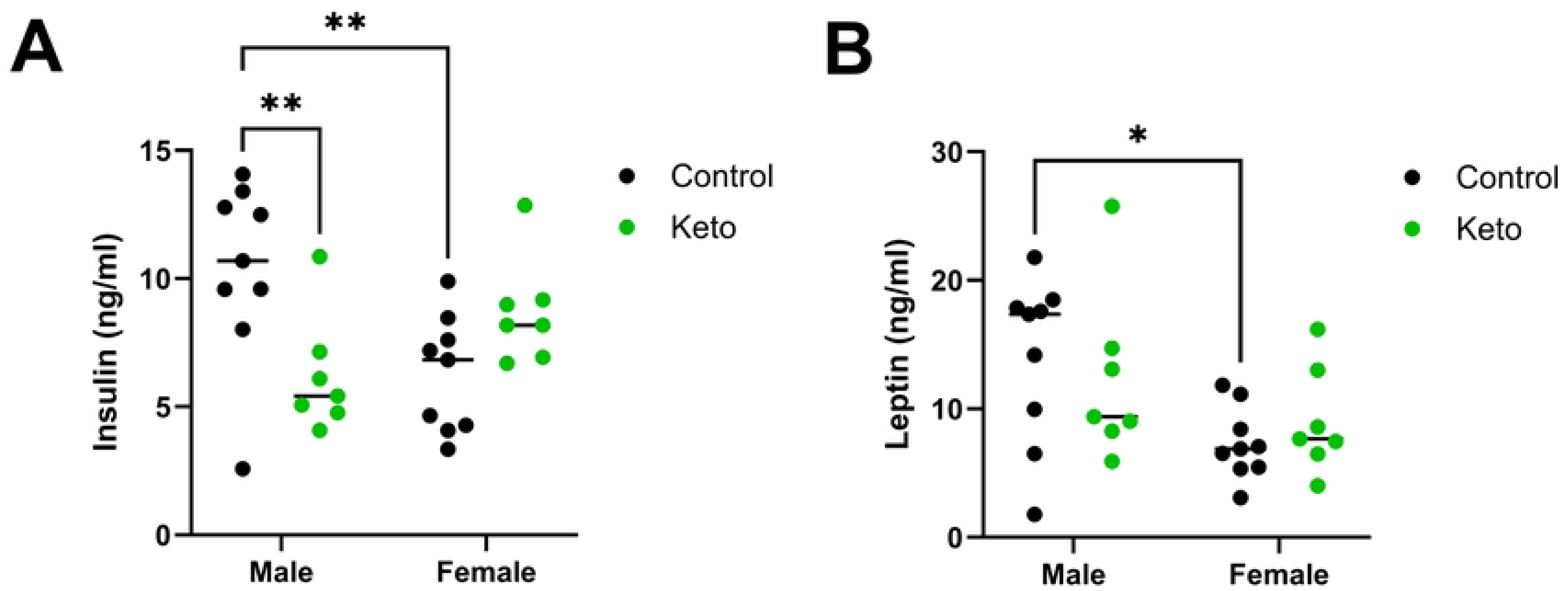
Figure 5**. Serum insulin and leptin levels in offspring at 12 weeks of age.** (A) Serum insulin concentrations. (B) Serum leptin concentrations. Data represent mean ± SE, with one male and one female per dam used for statistical analysis (Control n=9 dams, Keto n=7 dams). Statistical analysis was performed using two-way ANOVA with sex and maternal diet as factors. *p<0.05, **p<0.01.

Serum leptin levels did not differ between control and keto offspring (Fig. 5B). In control offspring, males exhibited higher leptin levels than females (p<0.05), whereas no sex differences were observed in keto offspring.

## DISCUSSION

The findings of this study indicate that maternal consumption of a ketogenic diet was associated with altered maternal glucose homeostasis during pregnancy and reduced fetal growth, as well as with modest alterations in offspring metabolic outcomes. Despite the limited maternal sample size, consistent differences were observed across multiple independent measures of maternal physiology, including glucose tolerance, urinary ketones, circulating insulin concentrations, maternal body weight, and fetal weight, providing internally consistent evidence that maternal ketogenic diet consumption altered maternal metabolism characterized by ketosis and glucose intolerance. However, key diagnostic criteria for diabetic ketoacidosis, including blood pH [32, 33], were not assessed, and therefore a definitive diagnosis of ketoacidosis cannot be made.

Data on the safety and metabolic consequences of ketogenic diet consumption during human pregnancy remain limited. A recent survey of healthcare professionals found that most providers do not recommend ketogenic diets during pregnancy, citing insufficient evidence regarding maternal and fetal safety [34]. In non-pregnant women, particularly those with PCOS, ketogenic diets are often associated with improved metabolic health and weight loss [17, 18]. In contrast, the ketogenic diet increased adiposity in our mouse model. This difference likely reflects species-specific metabolic adaptations and differences in diet composition. The ketogenic diet used in this study consisted of 75% fat derived from soybean oil, lard and cocoa butter and is comparable to diets used in prior rodent studies [21–23, 25]. Although murine models cannot fully replicate human metabolic response to ketogenic diets, they remine an important preclinical model for investigating the effects of maternal nutritional ketosis and carbohydrate restriction on pregnancy and offspring development under controlled experimental conditions. Consistent with our findings, Sussman et al reported gestational and lactational ketoacidosis leading to maternal mortality in mice [21], although no maternal deaths were observed in the present study, potentially due to differences in mouse strain or experimental conditions.

An alternative interpretation of the metabolic changes observed in the keto dams is that ketogenic diet consumption may augment the normal metabolic adaptations of pregnancy. Pregnancy is characterized by progressive insulin resistance and reduced peripheral glucose utilization, adaptations that promote glucose availability for the developing fetus [35, 36]. Under conditions of severe carbohydrate restriction, these physiological adaptations may be further exaggerated, resulting in altered glucose homeostasis rather than outright metabolic dysfunction. Additional studies assessing insulin sensitivity, circulating lipids, blood ketone concentrations, and whole body energy metabolism will be required to distinguish adaptive metabolic responses from pathologic metabolic impairment.

Although maternal metabolism was altered, offspring exhibited relatively modest metabolic alterations during early adulthood. Offspring exposed to a maternal ketogenic diet displayed reduced body weight during early development, consistent with the reduced fetal growth observed at GD17.5, but demonstrated catch-up growth by 12 weeks of age. While catch-up growth can normalize body size, it has been associated with increased risk of metabolic disease later in life, and longer-term studies will be necessary to determine whether adverse outcomes emerge with aging [37].

Interestingly, offspring from keto dams exhibited a modest improvement in glucose tolerance at 12 weeks of age. Although statistically significant, the magnitude of this effect was small, and its physiological relevance remains unclear. This finding suggests that early metabolic adaptations may occur in response to maternal ketogenic diet exposure, but do not necessarily indicate improved long-term metabolic health.

Sex-specific effects were also observed. In control offspring, males exhibited higher circulating insulin and leptin levels than females, consistent with known sex differences in metabolic regulation [28, 38, 39]. In contrast, these sex differences were attenuated or absent in offspring exposed to the ketogenic diet, suggesting that maternal diet may influence the development of sexually dimorphic metabolic pathways.

The reduced fetal weight observed in keto-exposed pregnancies despite comparable litter sizes suggests impaired fetal growth in response to maternal ketogenic diet consumption. However, because these findings are based on a limited number of pregnancies, confirmation in larger cohorts will be necessary. Reduced fetal size may be attributed to alterations in the intrauterine environment, potentially driven by maternal ketosis and impaired glucose metabolism. Ketone bodies are known to cross the placenta barrier and may also be present in breast milk, potentially exposing the developing fetus and neonate to altered nutrient availability [32, 40]. However, ketone concentrations were not directly measured in fetal or neonatal tissues in this study, and the mechanisms underlying fetal growth restriction remain to be determined.

It is important to note that offspring health cannot be assessed solely by the absence of early metabolic dysfunction. Previous studies have shown that maternal ketogenic exposure can adversely affect offspring neurodevelopment [20, 21]. Keto offspring have been reported to exhibit reduced white matter volume, impairing cognitive and motor development, likely due to altered lipid metabolism and elevated prenatal ketone exposure [19]. Other studies demonstrate reduced size of crucial midbrain structures alongside disproportionate enlargement of the hypothalamus and medulla [20], changes that may contribute to long-term neurological and physiological dysfunction.

This study has several limitations. Maternal and offspring outcomes were assessed in separate cohorts, precluding direct correlation analyses between maternal metabolic status and offspring phenotypes. Additionally, the sample size in the maternal cohort was limited, consisting of five control and three keto dams. Although statistically significant differences were observed across several independent maternal outcomes, the small sample size limits statistical power and increases the potential influence of biological variability. Therefore, these findings should be interpreted cautiously and confirmed in larger cohorts. Furthermore, litters were intentionally left undisturbed immediately after birth to minimize maternal stress and pup cannibalism. Consequently, individual pup body weights, maternal body weights during lactation, neonatal survival, and litter-level sex ratios were not recorded. Therefore, we cannot determine when the observed differences in offspring growth first emerged or distinguish prenatal from lactational influences of maternal ketogenic diet exposure. Future studies incorporating larger maternal cohorts, longitudinal assessments, and comprehensive metabolic assessments, including lipid metabolism, oxidative stress, and placental function, will be important to better define the impact of maternal ketogenic diet exposure on offspring health.

In conclusion, maternal ketogenic diet consumption during pregnancy was associated with altered maternal glucose homeostasis and reduced fetal growth in this mouse model, while producing modest and sex-specific metabolic changes in offspring during early adulthood. These findings highlight the need for further investigation into the long-term consequences of ketogenic diet exposure during pregnancy.

## ACKNOLEDGEMENTS

ChatGPT was used to generate figure 1 (experimental design) and edit this manuscript for clarity and conciseness. AI was not used to generate or analyze any data presented in the manuscript.

## AUTHOR CONTRIBUTIONS

A.H.R assisted with tissue collection, serum analysis, performed data collection and analysis as well as wrote the manuscript. R.A.K. and I.A.C. conceptualized the project, secured funding from the Department of Obstetrics and Gynecology Resident Research Fund and helped edit the manuscript. S.H.R. performed animal work and serum analysis, assisted with data collection and data analysis and edited the manuscript. M.J.T assisted with animal work and data collection and edited the manuscript. K.A.P. led the project, helped with project conceptualization, oversaw all animal work, data collection and data analysis. K.A.P assisted with writing the manuscript and made final edits.

## DATA AVAILABILITY STATEMENT

All data generated or analyzed during this study are included in this published article and its Supplementary Information files.

---

**Supplementary Table S1:** Diet Composition Breakdown as provided by Research Diets.

**Supplementary Table S2:** Reproductive Characteristics from Experiment 1 and 2

